# Genome-wide alignment-free phylogenetic distance estimation under a no strand-bias model

**DOI:** 10.1101/2021.11.10.468111

**Authors:** Metin Balaban, Nishat Anjum Bristy, Ahnaf Faisal, Md. Shamsuzzoha Bayzid, Siavash Mirarab

**Author notes:** These authors made equal contributions (Metin Balaban, Nishat Anjum Bristy).

## Abstract

While aligning sequences has been the dominant approach for determining homology prior to phylogenetic inference, alignment-free methods have much appeal in terms of simplifying the process of inference, especially when analyzing genome-wide data. Furthermore, alignment-free methods present the only option for some emerging forms of data, such as genome skims, which cannot be assembled. Despite the appeal, alignment-free methods have not been competitive with alignment-based methods in terms of accuracy. One limitation of alignment-free methods is that they typically rely on simplified models of sequence evolution such as Jukes-Cantor. It is possible to compute pairwise distances under more complex models by computing frequencies of base substitutions provided that these quantities can be estimated in the alignment-free setting. A particular limitation is that for many forms of genomewide data, which arguably present the best use case for alignment-free methods, the strand of DNA sequences is unknown. Under such conditions, the so-called no-strand bias models are the most complex models that can be used. Here, we show how to calculate distances under a no-strain bias restriction of the General Time Reversible (GTR) model called TK4 without relying on alignments. The method relies on replacing letters in the input sequences, and subsequent computation of Jaccard indices between k-mer sets. For the method to work on large genomes, we also need to compute the number of k-mer mismatches after replacement due to random chance as opposed to homology. We show in simulation that these alignment-free distances can be highly accurate when genomes evolve under the assumed models, and we examine the effectiveness of the method on real genomic data.

## I. Introduction

The dominant methodology used in phylogenetic inference is to assemble and align sequences and to use the alignments as input to phylogenetic inference. However, a large body of work also exists on alignment-free [1]–[7] and even assembly-free methods for inferring phylogenies [8]–[12]. While, for the most part, the alignment-free methods have not been as accurate as alignment-based methods [1], [6], they do provide several benefits and also enjoy emerging applications. The most obvious advantage is that alignments are difficult to infer and forgoing them would make the inference pipeline simpler. The challenges are further exacerbated when working with genome-wide data where long sequences and large-scale events such as rearrangements make alignment even more challenging [13]. There is, therefore, a hope that by skipping the alignment step, we can eliminate the errors [13] that are known to occur in the alignment step and impact phylogenetic accuracy [14]–[16]. Noting that alignment-free methods still need ways to consider homology, we do not hold this hope for alignment-free methods especially when applied to small rearrangement-free sequences (e.g., single genes). However, it seems possible that alignment-free methods could provide a reasonable trade-off between accuracy, running time, and complexity of analyses, especially for analyzing genomes, where alignment remains a challenging problem [8], [13].

The main advantage of alignment-free methods may come from situations where alignments are not even possible. In particular, genome skimming has emerged in recent years as a promising method of acquiring genome-wide data inexpensively [17] by generating short reads from across the genome at low coverage (e.g., 1X). While such data cannot be assembled, mapping them against a reference genome when available [18], [19] or analyzing them in an assembly-free fashion when references are not available are now possible [11], [20]–[23]. Given reads with low coverage, alignment is simply not possible, leaving us with alignment-free methods as the only option. Among assembly-free methods, many use *k*-mers to compute distances between all pairs of species and then use distancebased methods to infer the phylogeny. Recent *k*-mer-based methods include those that can work with both assembled and un-assembled data and model low coverage [9], [11], [22].

Despite their practical benefits in terms of simplicity and scalability, alignment-free methods have limitations of their own. One of these limitations is the reduced complexity of sequence evolution models employed. The majority of alignment-free methods rely on the simplest possible model of sequence evolution, Jukes-Cantor (JC) [24], which assumes equiprobable bases and base substitutions. In contrast, alignment-based methods use more complex models, such as the general time-reversible (GTR) [25] model paired with models of rate variation across sites and further partitioning data to allow changing model parameters. The reliance on JC is not an oversight by the research community. In the absence of alignments, it is more difficult to design methods for more complex sequence evolution models that need to estimate parameters related to relative rates of substitutions among bases. The difficulties are exacerbated by the fact that for unassembled and unaligned data, sequences can come from either of the two strands, making it difficult to calculate some parameters of complex models and impossible to compute others [26]. For example, while describing their *k*-mer-based method Skmer, Sarmashghi *et al*. [11] proposed a trick that they conjectured could be used in conjunction with the well-known LogDet technique [27] to compute distances under the GTR model from unassembled reads. However, this trick, which relied on the strategy of substituting all occurrences of each letter by another letter in input sequences, was never put to test.

In this paper, we note that the method proposed by Sarmashghi *et al*. fails in two ways. First, it fails to consider the fact that reads can be of either strain and given mixed-strand data, the trick proposed by Sarmashghi *et al*. will fail. We go on to show a more complex algorithm, inspired by the original technique of Sarmashghi *et al*., that can estimate all the parameters needed to compute distances for a time-reversible no strain-bias model called TK4 [28]. The second issue is that a fundamental assumption of many *k*-mer-based method, including Skmer, that matching *k*-mers can only appear by homology for a large enough *k*, can easily be violated after letter substitutions, especially for genomes with unbalanced base frequencies. Luckily, the expected number of random matches between two *k*-mers from two random genomes can be derived [29], and we go one step further and compute the expected (containment) Jaccard between two unrelated genomes (Lemma 1). Using these calculations, we can correct for the effect of non-homologous *k*-mer matches, thereby rescuing the letter substitution idea. Using analytical calculations and simulations, we show that using this technique to compute distances under the TK4 model can improve the accuracy of computed distances compared to JC, especially when the distances are high and deviations from the assumptions of the JC model are sufficiently high. We then use real data to demonstrate that the use of TK4 model improves the concordance of phylogenetic trees inferred from alignment-free methods and those inferred from alignment-based methods, indicating improved accuracy. We end by discussing the limitations of the method.

## II. Materials and Methods

### A. Background information

#### Evolutionary model

Suppose that we have two homologous DNA sequences 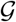 and 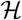 on character alphabet Σ = {*A, C, G, T*} taken from two species *S*_1_ and *S*_2_ that share a common ancestor. We assume that each homologous site in 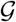 and 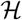 is evolved independently and according to a stationary continuous-time Markov-chain process on state set Σ that is defined by a 4×4 instantaneous rate matrix **R** = (*r_ij_*). Letting *π* = [*π_A_ π_C_ π_G_ π_T_*] denote the stationary base frequencies in 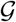 and 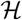, (thus, *π***R** = 0) the most general time-reversible stationary model, GTR [25], adds local balance constraints (i.e., ∀*i, j* : *π_i_r_ij_* = *π_j_r_ji_*), which lead to nine free parameters. Another constraint is added by requiring the time to be in the unit of one expected mutation, leaving us with eight free parameters. The transition matrix **P** = *e*^R*t*^ governs probabilities of base substitutions after time *t*.

Our goal is to estimate the time of divergence *t* between the two given genomes. Such estimates, if statistically unbiased, would converge to additivity and can be used with any distance-based phylogenetic inference method. In the last 50 years, numerous models with reduced complexity (i.e., fewer parameters) compared to the general Markov model have been proposed [24], [27], [30], [31], and some of these models have analytical equations for distance calculations [30], [31]. For example, let *genomic distance d* be the probability of observing a change in a homologous position. Under the simplest model, Jukes-Cantor (JC) [24], the maximum likelihood estimator is

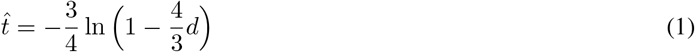

#### No-strain models

A restriction of GTR relevant to the study of NGS reads is the model proposed by Sueoka [32]. For a given base *i* ∈ Σ, let 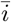 denote its complementary base (e.g., 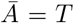). According to Sueoka’s no-strand-bias model (Fig. 1), an *i* → *j* substitution occurring on the forward DNA strand must have an identical rate to an *i* → *j* substitution occurring on the reverse DNA strand (parity rule 1). Since an *i* → *j* entails an 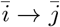 substitution on its opposite strand, the model requires the constraint 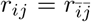 to hold, which reduces the number of independent parameters in the model to six. Sueoka postulated (despite other eveidence [33]) that when there is no selective pressure on one of the strands, data can be expected to manifest the model conditions [34].

**Fig. 1.**
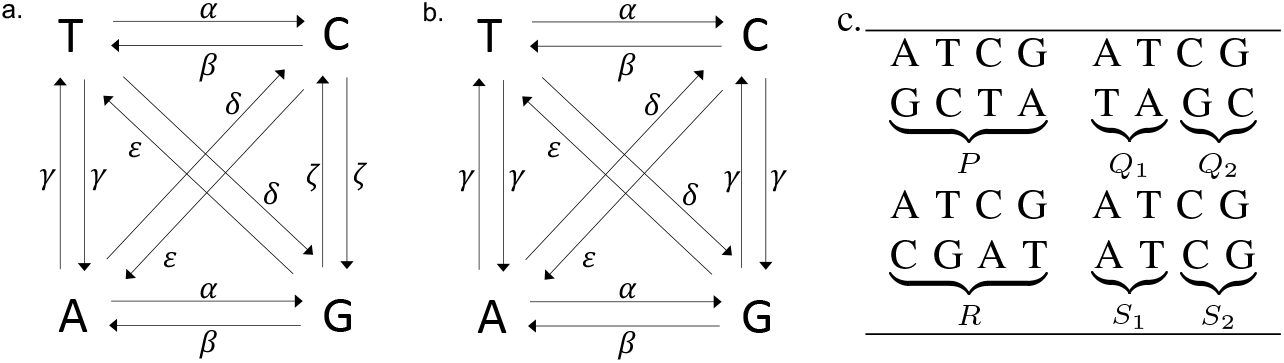
(a) Sueoka’s no strain-bias model of evolution with 6 rate parameters. TK5 model (b) is a special case of the 6-parameter model with the constraint *r_AT_* = *r_GC_*. TK4 is the time-reversible version of TK5 model with the condition 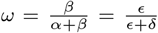 where *ω* is the total equilibrium frequency of bases *A* and *T*. (c) Nucleotide base pairs in homologous sites and their observed relative frequencies.

In this paper, we are dealing with conditions where the no strain-bias model is the *best* one can do. Assume 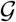 is not a single contiguous sequence but a set of *n* homologous sequences 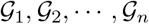 (similarly for 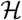) where each homologous pair 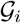 and 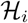 comes from an arbitrary strand. Such conditions arise when the input comprises *k*-mers, reads, or contigs. Under this assumption, *r_ij_* is unidentifiable from 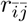 in an alignment-free pipeline. While the no-strand-bias model is useful, unfortunately, it does not allow analytical calculation of distance [26].

#### TK4

Predating Suoeka’s paper by 14 years, Takahata and Kimura introduced a 5-parameter non-time reversible model called TK5 [28] (Fig. 1b) that imposes on the general 6-parameter model the constraint *r_AT_* = *r_TA_* = *r_GC_* = *r_CG_* = *γ* and assumes that *π_A_* = *π_T_* = *ω*/2 and *π_C_* = *π_G_* = (1 − *ω*)/2. By imposing 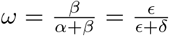, Takahata and Kimura introduce a time-reversible version of TK5 model with 4 parameters, called TK4 [28], and derive an analytical formula for distance estimation under TK4. This equation uses 16 combinations of bases possible at each site, as summarized in Figure 1c. Let *f_ij_* for *i, j* ∈ Σ denote the relative frequency of sites where the first and second genome has character *i* and *j*, respectively. For example, *P* is equal to *f_AG_* + *f_GA_* + *f_TC_* + *f_CT_*. Note that *P* + *Q*_1_ + *Q*_2_ + *R* + *S*_1_ + *S*_2_ = 1. An unbiased estimated phylogenetic distance 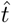 between 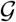 and 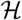 is given in terms of relative frequencies as follows^1^:

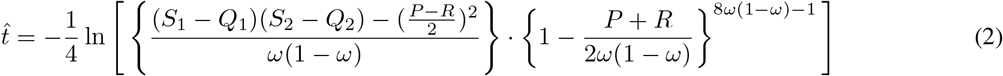

where *ω* can also be written as:

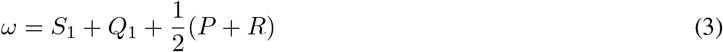

Comparing (1) and (2), it is not immediately obvious if the differences are consequential. By plotting the relative difference between (1) given the expected hamming distance under TK4 and the true time *t*, we can see that when parameters diverge from JC in biologically plausible ways, the often-used equation (1) can underestimate the true distance by more than 25% (Fig. 2). For example, with an AT-rich genome with *ω* = 0.75, setting *α* = 4 but keeping all other parameters equal to JC leads to 8% and 16% bias for true distances *t* = 0.25 and 0.5, respectively. As expected, bias is reduced when TK4 parameters are all close to 1 (i.e., JC assumption). Overall, it seems that high levels of bias correspond to cases where some of the relative rates diverge from others while base frequencies also diverge substantially from 25% (both of which are biologically plausible).

**Fig. 2.**
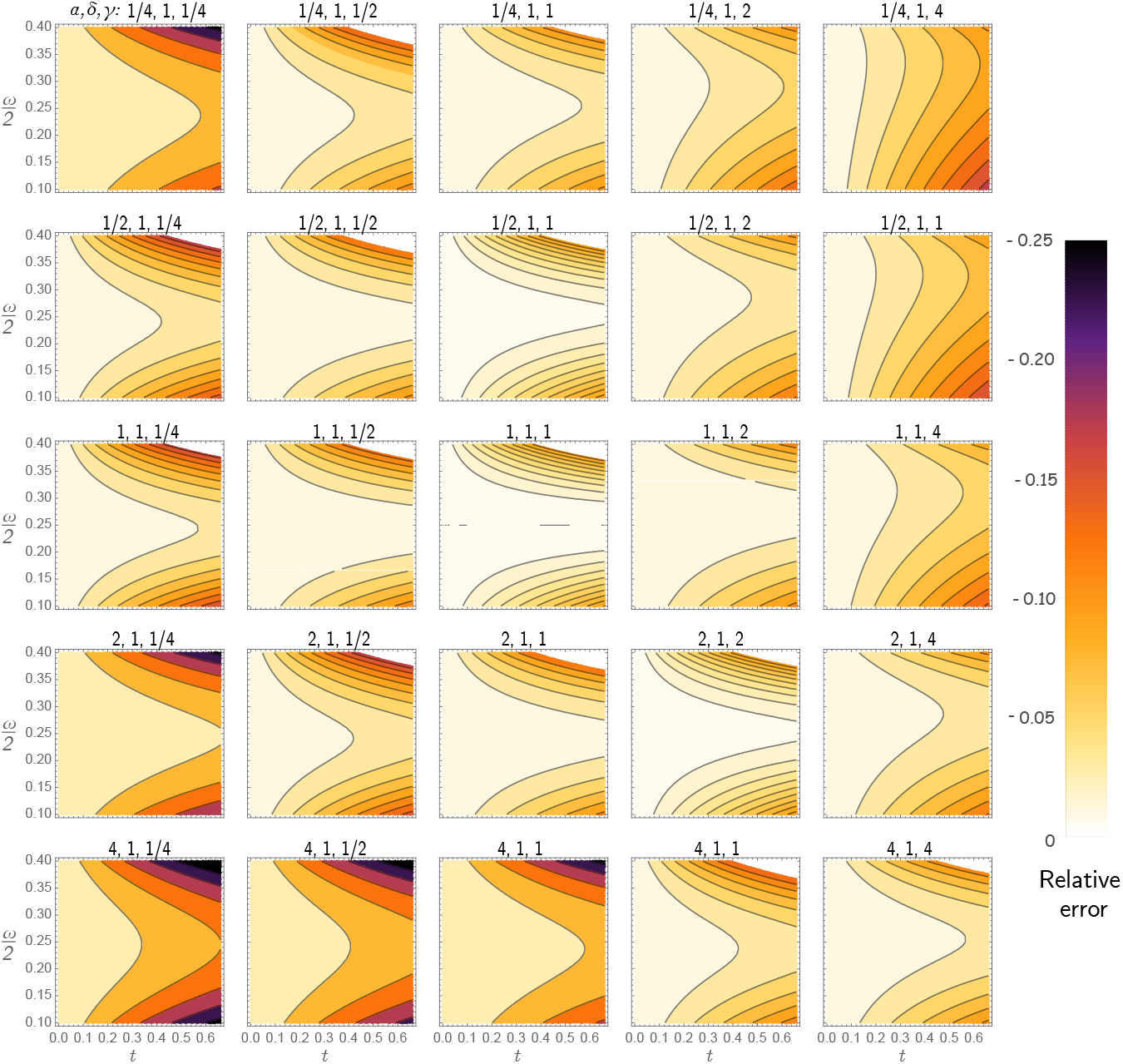
JC under-estimates TK4. The colors show relative bias of JC defined as 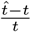 with 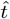 computed using equation (1) where *d* is set to the expected hamming distance under TK4, which can be computed as *d* = 1 − *π*.diag(*e*^**R***t*^). Each subplot corresponds to a choice of *α, δ, γ*, changing *α* across rows and *γ* across columns, fixing *δ* = 1. The x-axis changes the true evolutionary distance *t* and the y-axis changes the base frequency parameter *ω*. Note that the JC model corresponds to *α* = *δ* = *γ* = 4*ω* = 1.

#### Assembly-free distance estimation

Although it is trivial to compute observed frequencies of substitutions between two aligned sequences, such calculations are challenging in the absence of alignment, for instance when inputs are sets of unassembled reads. In the assembly-free setting, most methods assume the simple Jukes-Cantor [24] model (JC), which only requires genomic distance. Luckily, various alignment-free methods can estimate *d* [8], [11], [35], [36]. Many such methods [11], [35] break down the genome skims into *k*-mers.

We assume that a genome 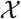 is a finite i.i.d. stochastic process *X*_1_*X*_2_…*X_L_* where each random variable (site) *X_m_* is drawn from categorical distribution with probability distribution *P*[*X_m_* = *A*] = *P*[*X_m_* = *T*] = *π_A_* = *π_T_* = *ω*/2 and *P*[*X_m_* = *C*] = *P*[*X_m_* = *G*] = *π_C_* = *π_G_* = (1 − *ω*)/2. A *k*-mer at position *m* is *X_m_X*_*m*+1_…*X*_*m*+*k*−1_ and denoted with *x_m_* in short. We denote the set of all *k*-mers in 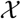 with 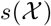. When *k* is sufficiently large with respect to *L* and *ω*, we can assume that 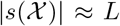. A second genome 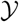 is originated from 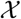 through a substitution process, described earlier. The probability of a match between two homologous *k*-mers is (1 − *d*)^*k*^. Therefore, the expected total number of homologous matches between 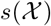 and 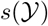 is approximately *S* = *L* · (1 − *d*)^*k*^ [9], [11], [35]. Denoting by 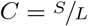 the containment Jaccard index, note that

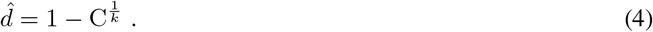

The *Jaccard index J*, defined as the intersection divided by the union of two sets, is easy to compute using techniques such as min-hash [35]. Thus, instead of *C*, most methods have relied on *J*, which is intimately connected to *C*:

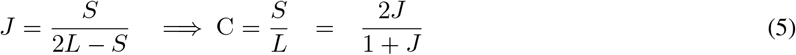

Finally, note that following the TK4 notations, 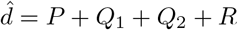.

### B. Containment Jaccard correction

In addition to homologous ones, *k*-mers in non-homologous positions in the two genomes can also match, albeit with lower probability. Distance estimation using Jaccard index requires computing the number of shared *k*-mers through homology. The number of non-homologous matches contributing to 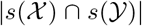 is negligible in most settings when *k* is large enough for the size of the alphabet; e.g., *k* = 31 with |Σ| = 4, leading to 4^31^ ≈ 4 × 10^18^ possible *k*-mers. However, as we will see, our algorithm for estimating TK4 distances requires reducing the alphabet set to three letters, and based on the value of *ω*, may lead to biased probabilities. Under such conditions, the non-homologous matches cannot be ignored.

Rohling *et al*. [29] have derived an expression for the expected number of *k*-mers *x_m_* and *y_n_*, *n* ≠ *m* that match between the two genomes by chance (i.e., not through homology). However, to compute the contribution of non-homologous matches to 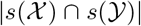, not only we need to know the expected number of *k*-mers matching by chance, we also need to account for a *k*-mer *x_m_* matching multiple *k*-mers in the other genome. Consequently, we propose a more precise estimate for the cardinality of intersection between two random genomes.

#### Lemma 1.

*The expected value of containment Jaccard for *k*-mers between two genomes* 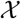 *and* 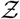 *generated by two i.i.d processes with stationary distribution π_A_* = *π_T_* = *ω*/2 *and π_C_* = *π_G_* = (1 − *ω*)/2 *is*:

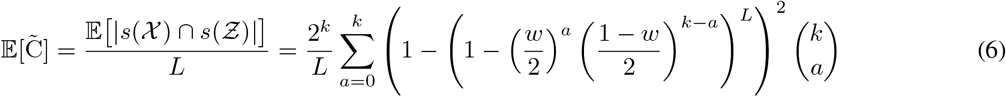

*Proof*. For 0 ≤ *a* ≤ *k*, let *r* ∈ Σ^*k*^ be a *k*-mer with *a* A’s and T’s.

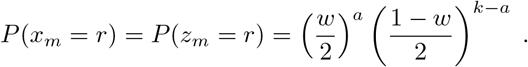

Therefore, the probability of *r* being in set 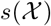 is the following

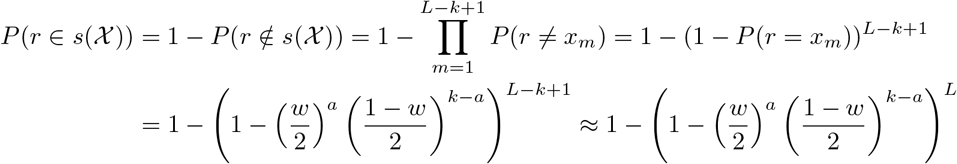

Results follows by noting that there are 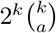 many selections for each *r* and 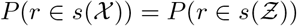.

By stationarity of the substitution process, 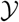 has the same base frequencies as 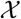. Thus, 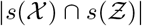 can be used to estimate the non-homologous portion of 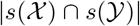. In other words, 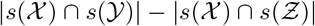 is the number of homologous *k*-mers. Combining Eqn. (4) and (6), *d* can be estimated from the observed containment Jaccard *C* between 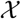 and 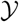:

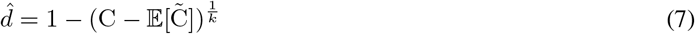

### C. Calculation of TK4 terms via replacement

Given the possibility of high error with the JC model (Fig. 2), we would like to develop alignment-free methods of computing distances according to the TK4 model using (2). Therefore, our goal is to *estimate* the terms *P, Q*_1_, *Q*_2_, *R*, *S*_1_, *S*_2_, and *ω*. Consider the replacement technique where every occurrence of a character *i* ∈ Σ in 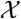 and 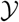 is replaced with character *j* ∈ Σ, *i* ≠ *j*. Let *d_ij_* be the genomic distance between two genomes after such replacement. The reduction in genomic distance after *i* to *j* substitution is exactly *f_ij_* + *f_ji_* Using the Eqn. (7), *d_ij_* can be estimated from empirical containment Jaccard *C_ij_* and expected number of background matches 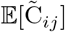. Using this replacement scheme, the *P*, *Q*_1_, *Q*_2_ and *R* terms in (2) are estimated as follows:

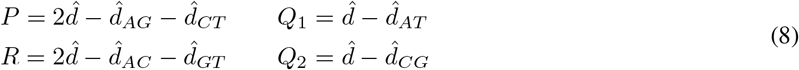

As base frequencies 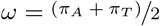 can be trivially computed from 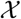 and 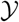, we can compute the remaining terms *S*_1_ and *S*_2_ using (3):

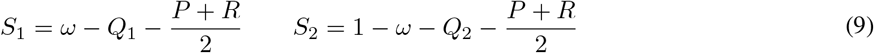

As mentioned previously, estimating *d_ij_* requires computation of 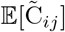. Calculation of this term depends on the type of replacement. Lemma 1 can be easily updated to account for replacements. For instance,

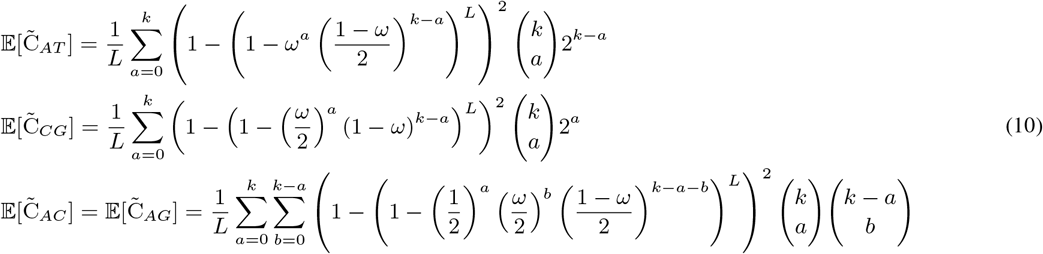

Since letter replacements, especially A to T for *ω* > 0.5 and G to C for *ω* < 0.5, can lead to a very high expected number of shared *k*-mers by chance, the use of these equations to correct for their effect is essential. For example, with a pair genomes of length 10^8^ and *ω* = 0.6, the expected number of background matches between 2-way genomes after A-to-T replacement is 289,000, which is 5× larger than the number of homologous matches when *t* = 0.5. Figure S1 shows the accuracy of equations (10) and their improvement over simply using the expected number of *k*-mer matches, as derived by Rohling *et al*. [29].

### D. Handling mixed-strand conditions

We now consider the case in which each k-mer in 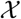 and 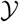 may come from forward or reverse DNA strand arbitrarily. In practice, chromosomes or contigs in an assembly, or reads in a sequencing run may come arbitrarily from either one of the forward or reverse strands. For simpler exposition, assume each genome consists of a single contig from an unknown strand (no such assumption needed by method). Let 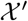 be another finite i.i.d. stochastic process 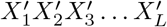 such that is 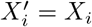 with some unknown but fixed probability *p_x_* > 0 and 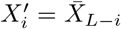 with probability 1 − *p_x_* where 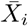 is reverse complement (RC) of *X_i_*. 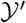 is defined similarly. Genomic distance between 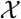 and 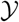 can still be computed using (4) by using canonical k-mers, a concept utilized by several tools [35], [37]. We utilize the same concept and construct a 2-way genome 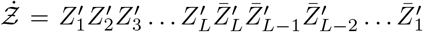 with 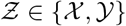 by adding the RC of each genome to itself. By design, both forward and reverse copies of each k-mer in 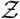 are present in 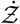. If *x_m_* = *y_m_*, either 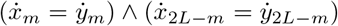 or 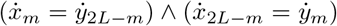. Either way, the number of homologous matches and genome length both double compared to the case where all sequences are of the same strand, leaving containment Jaccard due to homologous *k*-mers unchanged; thus, Eqn. (7) is applicable to 2-way genomes with the only change being that 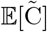 should be computed with 2*L*.

Similarly to the replacement technique shown previously, we introduce *i* to *j* replacements on a 2-way genome. For each homologous site (*X_m_, Y_m_*) in the base genomes 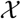 and 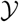, we have two pairs of homologous sites in 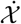 and 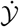. Although there are 4 alternative choices for assignment of forward and reverse strain to 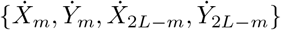, without loss of generality, let 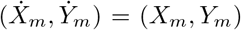 and 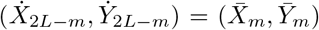. After replacing every occurrence of *i* ∈ Σ with *j* ∈ Σ in 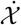 and 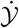,

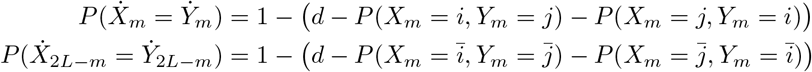

The reduction in genomic distance between 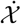 and 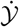 after the replacement, 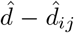, is *f_ij_* + *f_ji_* in the forward strand (i.e. 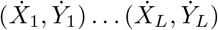) and 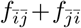 in the reverse strand (i.e. 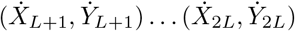). The overall reduction is the average of the reduction in the forward and reverse strands, which is 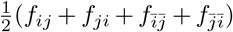. As a result, 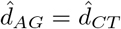 and 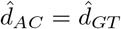. The *P, Q*_1_, *Q*_2_ and *R* terms in (2) are estimated from 2-way genome using:

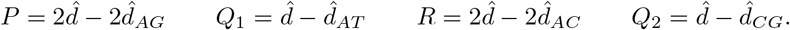

Thus, our estimation procedure requires computing only five distance values from the data, 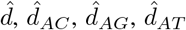, and 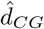 in addition to an estimation of *ω*.

Although four TK4 terms *P, Q*_1_, *Q*_2_, and *R* can be determined independently given the estimate 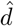, they must satisfy the constraint 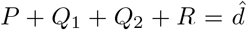. Thus, 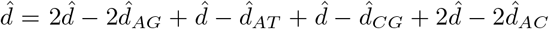. Since all five estimated values 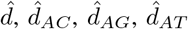, and 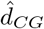 are empirical, it cannot be ensured that this equation will be satisfied. In other words, the system of equations has one excess observation. Among the five, the distance with no replacements 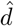 is always the largest, i.e. has the lowest containment Jaccard index. For large distances, the containment Jaccard can be zero, which prohibits computing any evolutionary distance (JC or TK4) from the data. In order to increase the distance upper-bound of *TK*4 model, we opt to reduce the number of free variables in the system by computing 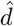 from 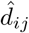, not directly from data. More precisely,

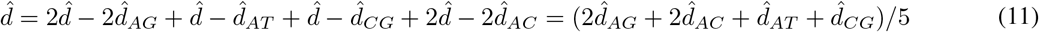

This equation can be used to compute JC model distances using (1). We also use it to calculate *P*, *Q*_1_, *Q*_1_, and *R* as a linear combination of four 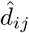 distances calculated using (7) after replacement:

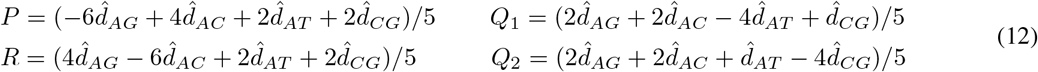

### E. NSB: An algorithm for TK4 distance estimation using k-mers

Algorithm 1 combines results in the previous sections into a three-step process (Fig. 3) for estimating phylogenetic distances under the TK4 model. We implemented the algorithm using Python in a method called the NSB (No Strand Bias) distance estimator. In its first step, NSB adds the reverse complement of all input sequences. It then builds separate *k*-mer libraries for each of the inputs using a left/right encoding scheme where nucleotide bases *A*, *C*, *G* and *T* are represented as 2-bit numbers, thus requiring 64-bit integer for *k* <= 32. NSB then builds base substituted encoded *k*-mer libraries from the initial encoded library by replacing the encoded bits of base *i* with the encoded bits of base *j*, for (*i, j*) ∈ {(*A, C*), (*A, G*), (*A, T*), (*C, G*)}. Thanks to a Left/Right encoding scheme, a replacement operation on an array of *k*-mers can be computed rapidly using fast and vectorized bitwise operations such as XOR, AND, and Shift (e.g., see A_to_C function in Algorithm 1). Finally, NSB computes the Jaccard indices for 4 pairs of base-substituted encoded libraries by computing the cardinality of the intersection succeeded by containment Jaccard correction. In practice, input genomes are almost never the same size and never follow the same base frequencies. When computing 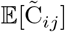 using Lemma 1, 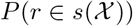 and 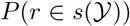 are computed using *L* and *ω* of the respective genome for a given *k*-mer *r*. In the final stage, we estimate the phylogenetic distance of each pair of genomes using Equation (2). Various components in this equation are calculated using the equations (9)–(12). *L* and *ω* are set to the average of the two input genomes. This procedure has *O*(*nN* + *n*^2^*N* log(*N*)) running time and *O*(*N*) byte of memory for calculating all pairwise TK4 distances for a set of *n* genomes of length *N* (Algorithm 1.)

**Fig. 3.**
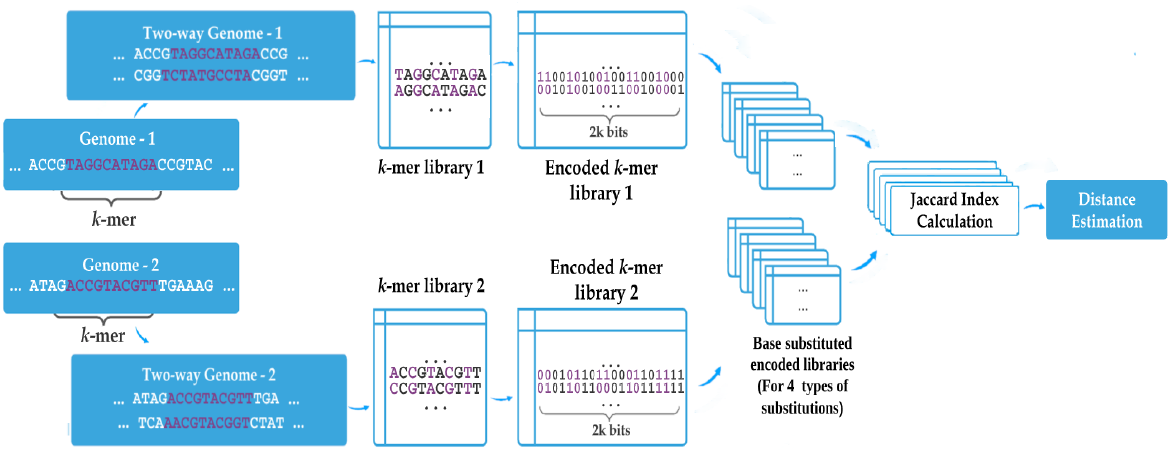
Schematic diagram of our proposed pipeline. We start with adding RC of sequences followed by decomposing them into fixed-length *k*-mers. Next, the bases in *k*-mers are encoded as bitmasks and thus an encoded *k*-mer library is constructed. For each (*i, j*) ∈ {(*A, C*), (*A, G*), (*A, T*), (*C, G*)}, we replace the encoded bits of base *i* with the encoded bits of *j*, producing 4 base-substituted encoded libraries. Finally, using these encoded libraries, Jaccard indices and distances are estimated under the assumptions of the TK4 model and using the equations derived and presented in Sec. II.

#### Algorithm 1 NSB: TK4 Distance estimation.

Notations: We denote the set of all reference sequences by *S*. NSB first runs PREPROCESS procedure, which itself uses ADD_RC to add the RC of genomes. It then calculates pairwise distances of the sequences according to the PAIRWISE-DIST procedure. BG_INTERSECT computes expected number of background matches after replacement the using equation (10).

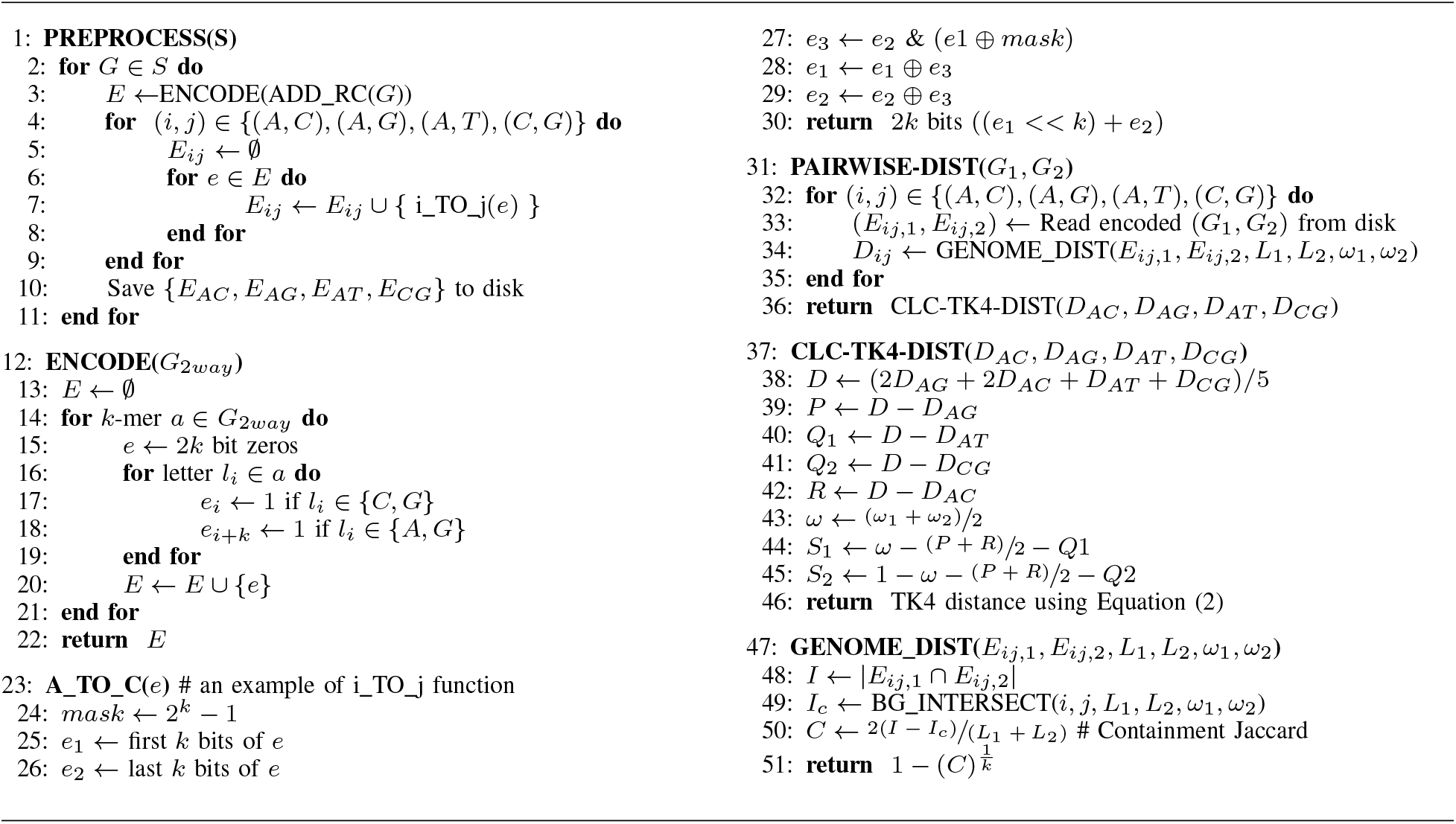

## III. Validation and benchmarking

We next validate NSB in simulation studies and on real data. We compare NSB-TK4 to three other methods. NSB-JC is Jukes-Cantor (JC) distance computed using (11) and (1) with our tool. We also test two existing tools, Jellyfish and Skmer, to estimate (containment) Jaccard index and subsequently JC distance using (1) and (7). While Jellyfish computes the Jaccard index exactly, Skmer distances are approximations based on 10^5^ sketches.

### A. Simulation study

#### Simulation process

We use a simple procedure to simulate pairs of genomes evolved under the TK4 model with controlled levels of distance and model parameters (https://github.com/balabanmetin/tk4-evol-sim). First, we either use a real genome as the ancestral genome or simulate one by drawing each site randomly from *π* with user-defined *ω*. We simulate two separate genomes from the ancestral genome by introducing substitutions at random positions. The frequency of each substitution type is determined by the TK4 model transition probability matrix **P** and half of the targeted distance *t*/2, producing two genomes with the evolutionary distance *t*. We create two simulated datasets. The first dataset uses a randomly generated 100*Mbp* base genome with *ω* = 0.6. The second dataset uses a real assembled genome of *Saccharomyces arboricola* (11 Mbp) as the base sequence. The base frequencies of the available *S. arboricola* genome are *π_A_* ≈ *π_T_* ≈ 0.307 and *π_C_* ≈ *π_G_* ≈ 0.193, which follow the assumptions of TK4 with *ω* = 0.614. We set the parameters of the TK4 model according to Fig. 1, exploring eight values of *α, δ*, and *γ*. Recall that 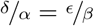 and 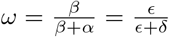, leaving us with only three free parameters for a fixed *w*. We generated eight model conditions with different TK4 parameters (Table S1) chosen to include cases with both minimal and substantial deviations from the JC model based on the earlier calculations (Fig. 2). For each model condition, we simulated genome sequences with true distances *t* ∈ {0.01, 0.05, 0.1, 0.2, 0.3, 0.4, 0.5}, each with 10 replicates, covering a range of both short and long distances.

#### Results on simulated datasets

##### a) Random base genomes

When input genomes are generated in the i.i.d. fashion assumed by both evolutionary models, across all model conditions and regardless of the true phylogenetic distances *t*, the distances estimated by NSB-TK4 are highly accurate (Fig. 4). In contrast, JC distances are accurate when the true distance *t* is low but are under-estimated when *t* increases. In the most challenging case, *t* = 0.5, NSB-TK4 deviates only 0.3% from the true value on average compared to 7.8% for Jellyfish-JC. The percent deviation of Jellyfish-JC is as high as 18% when *γ* = 32, causing extreme deviations from JC. The best performance of JC is when all parameters except *ω* follow JC. As models become successively more deviant from JC assumptions, the accuracy of JC diminishes.

**Fig. 4.**
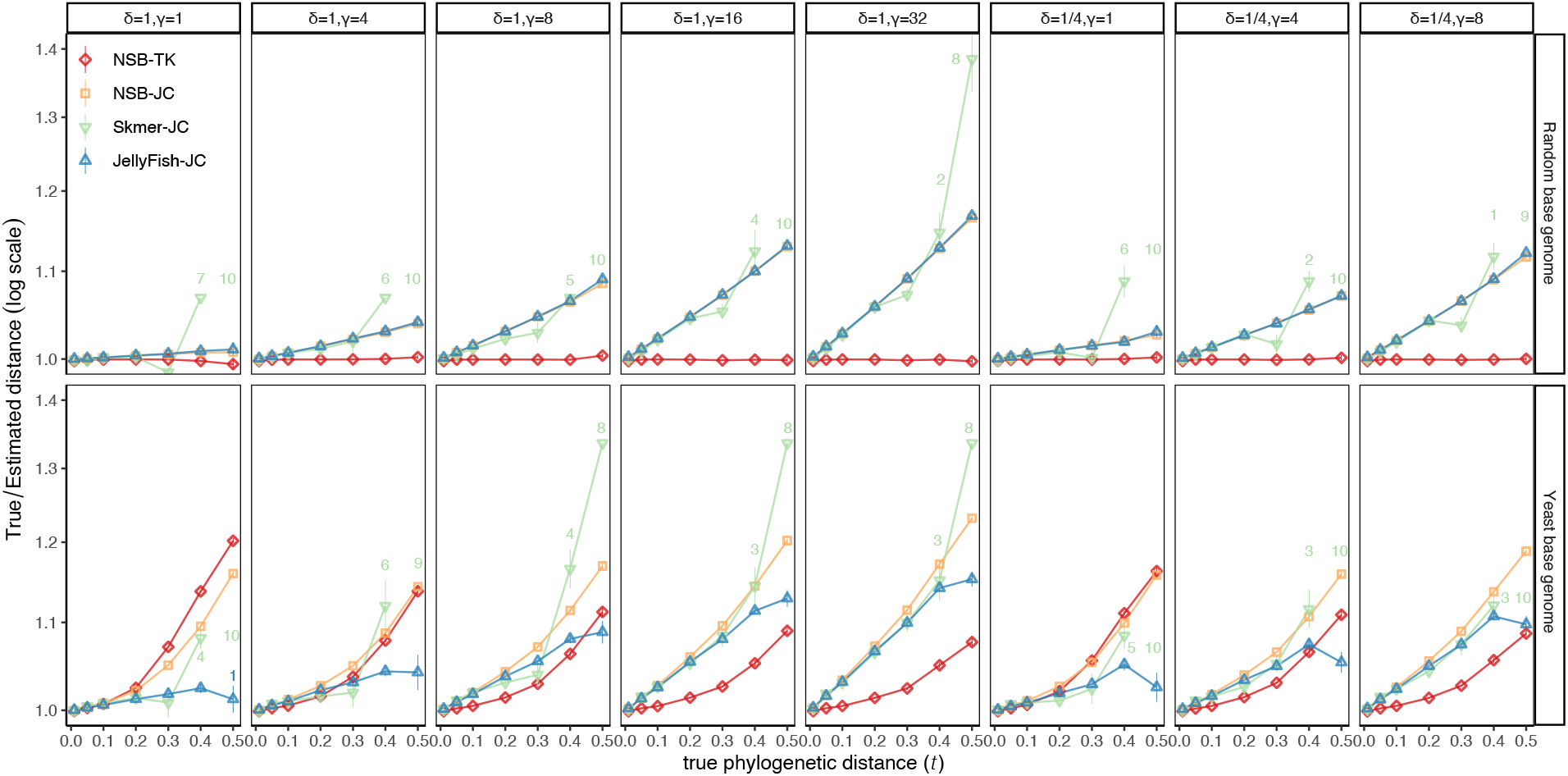
Comparing the accuracy of distances estimated by different approaches on random and Yeast-based simulated genomes. Genome sequences were simulated by randomly substituting the genome skims of *Saccharomyces arboricola* (11 Mbp) and a random 100 Mbp sequence with eight different sets of TK4 parameters and with seven controlled true distances. Here, *ω* is fixed, and since these rates do not have a scale, *α* =1 in all cases. We show the average true distance divided by estimated distances (y-axis) with standard errors (over all replicates, requiring at least two) against the true distances. Annotated numbers show the number of replicates out of 10 where Skmer or JellyFish return infinity.

Finally, comparing the two ways of obtaining JC distances, for *t* ≤ 0.3, surprisingly, the approximate Skmer distances are slightly *more* accurate than Jellyfish. However, when *t* > 0.3, Skmer distances become less accurate. When true distance *t* ≥ 0.4, Skmer fails to estimate distances in some cases (most cases for *t* = 0.5) because the true Jaccard index becomes too small (e.g., < 10^-5^) to compute reliably with sketches of size 10^5^.

##### b) Yeast-base simulations

The TK4-based calculations show improvements over JC computed using NSB or Skmer across some model conditions except for *δ* = *γ* = 1 that resembles JC (Fig. 4). However, the comparison to JC computed exactly (using JellyFish) is more complex. When deviations from JC are relatively low, JellyFish-JC can be as accurate or even more accurate than NSB-TK4. It is only with higher levels of deviation from JC that improvements of NSB-TK4 over JC are clear. Regardless of simulation parameters, phylogenetic distances *t* ≤ 0.1 are estimated with high accuracy under both TK4 and JC models. However, the JC model starts to underestimate the distance as we increase the distance *t*, and the underestimations are substantial when *t* ≥ 0.3. Moreover, the JC error is not linear or even monotonically increasing with respect to *t*, meaning that the distance matrices obtained from the JC model may not be additive. When *t* is increased to 0.5, TK4-based distances tend to have reasonable accuracy with a few exceptions (e.g., for *γ* = 8). In some cases, TK4 distances have more than 10%error with increased *t* and in three conditions are consistently less accurate than JC. Comparing the results to random base genomes, the reduced accuracy of TK4 on these conditions has to be due to violations of the model in the base genome, a point that we will return to in the discussion section.

### B. Evaluation on biological bacterial data

#### Dataset

We created a dataset consisting of 10 clades of microbial species subsampled from the Web of Life (WoL) [38] ASTRAL tree of 10575 Bacteria and Archaea taxa. We started with finding all the clades with 30 to 50 leaves and with 0.2 to 0.7 diameter (the largest pairwise tree distance between any pair). We then selected the top 25 clades with the highest local posterior probabilities and for each clade, computed an all-pairwise distance matrix using Skmer (sketch size 10 million), inferred a phylogenetic tree using FastME 2.0, and computed the Robinson-Foulds (RF) [39] distance between the WoL ASTRAL reference tree and the inferred tree. We then selected nine clades with the lowest RF distance, and these clades had 32 to 46 species and RF distance between 0.16 and 0.42. As none of the nine selected clades had any missing data in their distance matrix, we also curated a challenging subtree with 86 taxa from the *Erysipelotrichaceae* family from the WoL reference tree that contained 114 missing data entries in its distance matrix (RF distance: 0.43) computed using Skmer.

#### Results on Bacterial dataset

On the 10 bacterial data sets, while methods are generally competitive (Fig. 5a), overall, NSB-TK4 is better than others as it produces the best result in 8 datasets out of 10. The total number of missing branches for NSB is 120 (out of 403) (Table S2), which is lower than Jellyfish, with 133 missing branches. Results are similar when focusing on highly supported branches: NSB-TK4 misses 95 out of 374 branches with at least 0.95 support while Jellyfish misses 109. Among the three methods that compute JC distances, NSB-JC is the most accurate, matching or improving on Jellyfish and Skmer in seven out of 10 cases, and with eight and four fewer wrong branches, respectively. On the most challenging case (Set 10), the distance matrix produced by NSB-TK4 contains 20 fewer missing entries (infinity) than both Jellyfish-JC and Skmer-JC. As a result of its replacement technique, NSB can compute distances where other tools cannot. To perform a tree inference on distance matrices with missing data, we impute the missing distances using a machine-learning-based algorithm [40]. Here, NSB-TK4 distances produce the tree with the fewest differences to the reference phylogeny compared to JC-based tools.

**Fig. 5.**
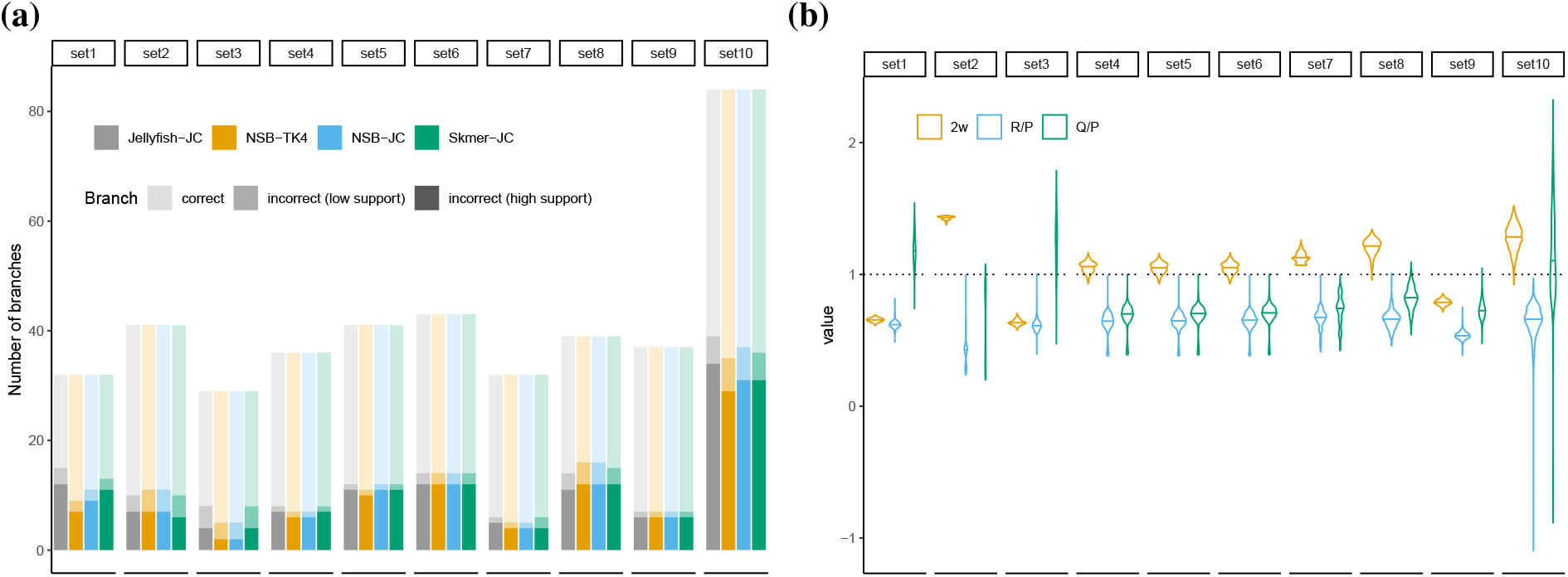
(a) Comparison of different methods to the ASTRAL tree on 10 subsets of the bacterial dataset. We show the number of branches in the reference tree that are correctly estimated, or are incorrectly estimated and have low (less than 0.95) or high support in the reference tree. (b) TK4 model parameters inferred using NSB-TK4 for each set. Deviations from *y* = 1 indicate violation of the JC model. *Q* = *Q*_1_ + *Q*_2_.

Beyond topological accuracy, we quantify the divergence of the TK4 and JC distances from the additivity using the Fitch-Margoliash [41] (FME) weighted least squares error. Since FME metric weights distances by 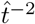, it is insensitive to the unit and scale of branch lengths. Jellyfish-JC had between 7% and 57% (mean: 22%) higher FM error than NSB-TK4 across datasets (Table. S3). NSB-TK4 distances are not only more additive but also on average 13%and 32%larger than those of Jellyfish and Skmer, which may under-estimate the distances.

TK4 model parameters inferred by NSB-TK4 demonstrate that JC model assumptions are significantly violated in the real data (Fig. 5b). For instance, 2ω, which is assumed to be 1 in the JC model, is as low as 0.65 on average across all pairs in a set. In addition, transversion to transition ratios *R/P* and (*Q*_1_ + *Q*_2_)/*P* are less than 1 in almost every case, as expected and in clear violation of the JC model, a problem that has been long understood [42].

### C. Evaluation on biological Yeast dataset

#### Yeast dataset

We also study a dataset with eight yeast assemblies from a previous study [21], consisting of eight genomes (Table. S4) with sizes varying between 10.9Mbp to 12.4Mbp. The number of chromosomes/scaffolds in the genomes highly vary between 16 and 2808. We use a previously published phylogeny for Yeast [43] as a reference, and compute it to alignment-free trees inferred under TK4 and JC models using FastME 2.0 [44].

#### Results on Yeast dataset

For the Saccharomyces species, our method NSB-TK4 and Jellyfish-JC produce a phylogenetic tree that is identical to the reference phylogeny (Fig. S2). However, Skmer-JC distances produce a tree with one branch mismatch. Although the trees inferred using NSB-TK4 and Jellyfish-JC distances are topologically identical, their branch lengths differ: NSB-TK4 trees have 16% increased tree height (Fig. S3), indicating that JC model likely under-estimates distances. In terms of additivity, Jellyfish-JC distances have an FME of 0.0034, which is 66% higher than that of NSB-JC.

## IV. Discussion

In this paper, we introduced an algorithm and a tool for computing phylogenetic distances on alignment-free data based on a time-reversible, no strain-bias, 4-parameter evolutionary model called TK4. Through theoretical and empirical analyses, we explored the model conditions where a general model like TK4 can offer more accurate distances than Jukes-Cantor model, which is the simplest yet most dominantly used model in alignment-free phylogenetics. As expected, the improvements are most pronounced for larger distances and for higher levels of deviations from the JC model assumptions.

Despite overall improvements, on the simulations based on the real yeast genome, we observed conditions where the TK4 model was less accurate than the JC model it contains. Deviations from the TK4 model may be behind this surprising result. The real biological data, even if used as the base genome for subsequent simulations, can violate the assumptions of our algorithm in several ways. *i)* Presence of non-randomly generated repeats (e.g., recent gene duplications) causes overestimating the Jaccard index. The probability of a *k*-mer to be present in both input genomes is higher when it repeats multiple times across the genome. Our calculations only correct for these repeats when they occur randomly but not by homology. *ii)* Systematic variations of *ω* across the genomes, violating i.i.d. assumptions, can create loci with increased numbers of homologous and non-homologous matches after replacement. *iii)* Presence of *k*-mer motifs can invalidate assumptions of Lemma 1. While some of these issues also violate JC assumptions, NSB-TK4 may be less robust to these violations than JellyFish-JC due to the more complex equation used for TK4 (2) or the more complex estimation procedure (including letter replacement) used by NSB.

More broadly, while the TK4 model is more complex than JC, important processes are also missed by TK4. An important aspect of molecular evolution that is not modeled in this study is the rate heterogeneity among sites. Leading alignment-based phylogenetic estimation tools offer modeling the heterogeneity using a discrete or continuous gamma distribution. JC model can be extended to support Gamma-distributed rates [45] if the parameters of the Gamma model are known. It may be possible to incorporate a measure of rate variation in the TK4 formula (2) as well. We leave that question to future work.

The method we described simply needs computing the (containment) Jaccard index between *k*-mer sets of the input genomes and their perturbations. This approach, previously utilized by tools such as Mash [35] and Skmer [11] for computing JC distances, has the potential to be applied to both assembled genomes and NGS reads in an assembly-free fashion. Nevertheless, applications directly on NGS reads also require dealing with sequencing errors and potentially a lack of coverage across the entire genome. Such corrections are previously introduced [11] and can be applied here, with the added complexity that random non-homologous matches need to be considered. While such analyses are possible, we leave the incorporation of these two parameters into NSB to future work.

The use of *k*-mers is not the only option for distance calculations. For example, tools like pyANI [46] and Co-phylog [8] estimate distances between two genomic sequences by efficiently finding local alignments. It is possible to infer substitution probabilities from these local alignments and calculate evolutionary distance according to the TK4 model. While such approaches will not be fully alignment-free, future work should compare these methods to our proposed approach. However, even if accurate, such methods cannot be incorporated into the analyses of low-coverage short-read NGS data mentioned above when assembly is not possible.

We used *k* = 31 in all benchmarks. However, previous results indicate that *k*-mer size may affect the accuracy in some alignment-free distance-based applications and have proposed a strategy for automated selection of the best *k* value [21]. Similarly, for the *k*-mer-based alignment-free distance estimation problem, techniques that enable an automatic choice of *k*, as used in other areas [47], may improve results.

Due to the exact computation of *k*-mer counts, NSB and JellyFish can both have substantial running times. On two random genomes of length 100Mbp, NSB completes within 15 minutes where 13 minutes is spent for preprocessing the samples and computing the encodings and less than 2 minutes for computing all 4 Jaccard values and the pairwise TK4 distance. Jaccard indices can be estimated accurately without looking at all *k*-mers using the MinHash sketching technique [35] that dramatically improves the running time, disk space, and memory usage. However, we saw that for large distances where Jaccard is small, MinHash sketching fails. This limitation may be alleviated with newer methods such as Dashing [48]. Nevertheless, for smaller distances where it is accurate, we could incorporate sketching into NSB. In preliminary tests, we saw that while the main Jaccard index is often computed accurately using sketching, the replaced Jaccard indices can have consequential levels of error. This is likely because hash functions used in existing tools assume four letters, and need to be updated for genomes with replaced letters. It may even be possible to compute all four Jaccard indices without actually replacing letters by defining hash functions that do not distinguish letters. Finally, NSB may be able to use compressed *k*-mer sets [49] to reduce its storage requirements while keeping the same accuracy. We leave the exploration of these avenues to further work.

## V. Availability

Our software is available open source at https://github.com/nishatbristy007/NSB. The Yeast dataset is available at https://github.com/balabanmetin/yeast-genomes. The bacteria dataset is available at https://github.com/balabanmetin/bac10.

## Acknowledgment

This work was supported by the National Science Foundation (NSF) grants [NSF-1815485 to M.B. and S.M.].

## S1. Supplementary Figures

**Fig. S1.**
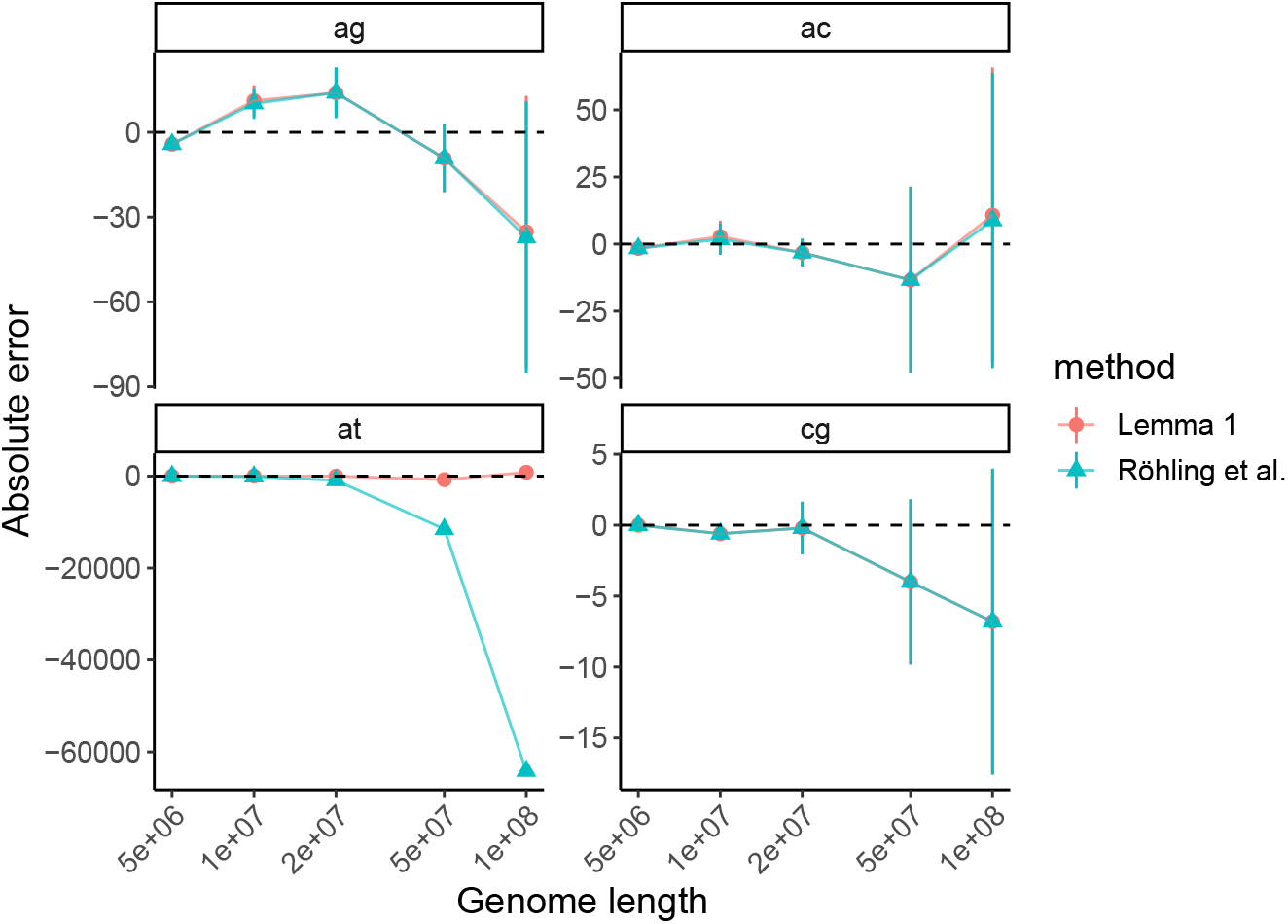
Estimating number of non-homologous (by random chance) matches between two genomes after replacements. Two input genomes are created according to an i.i.d. process with base frequencies *π* = [*π_A_* = 0.3 *π_C_* = 0.2 *π_G_* = 0.2 *π_T_* = 0.3]. Replacements are performed after 2-way genomes are constructed.

**Fig. S2.**
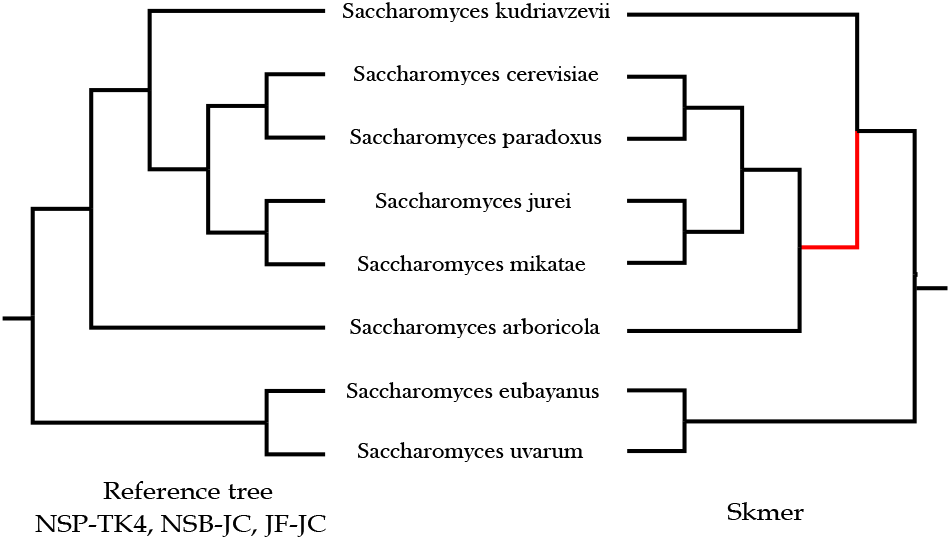
Analysis of the yeast genomes using distances estimated under JC and TK4 models. The branch in the estimated tree which is not found in the reference tree is shown in red.

**Fig. S3.**
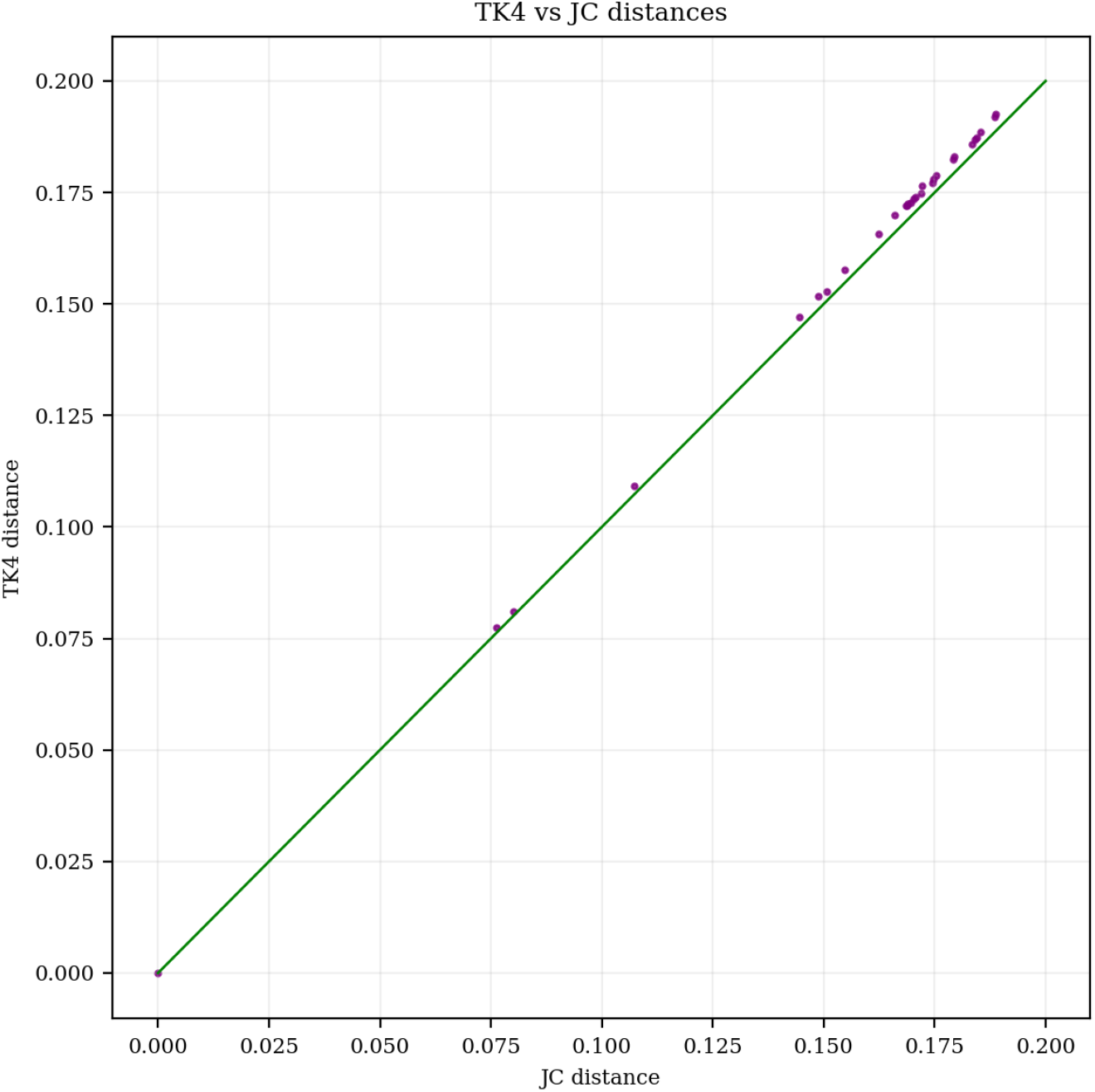
Comparison of pairwise distances estimated using TK4 and JC models on 8 real Yeast genomes. TK4 distances are always slightly higher than JC.

## S2. Supplementary Tables

**Table s1.**
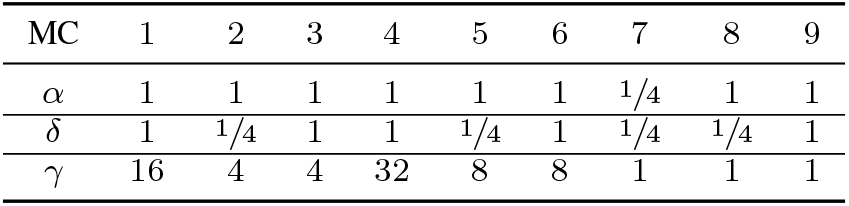
Different model conditions (MC), used for simulating genome sequences

**Table s2.** Comparison of different methods to the ASTRAL tree on 10 sets of bacterial dataset. Best results are shown in bold.

|  | # taxa (branches) | Number of branch mismatches with ASTRAL |  |  |  |
| --- | --- | --- | --- | --- | --- |
|  |  | NSB-TK4 | NSB-JC | Jellyfish-JC | Skmer-JC |
| Set 1 | 34(31) | **9** | 11 | 15 | 13 |
| Set 2 | 43(40) | 11 | 11 | **10** | **10** |
| Set 3 | 32(29) | **5** | **5** | 8 | 8 |
| Set 4 | 38(35) | **7** | **7** | 8 | 8 |
| Set 5 | 43(40) | **11** | 12 | 12 | 12 |
| Set 6 | 46(40) | **14** | **14** | **14** | **14** |
| Set 7 | 34(31) | **5** | **5** | 6 | 6 |
| Set 8 | 41(38) | 16 | 16 | **14** | 15 |
| Set 9 | 39(36) | **7** | **7** | **7** | **7** |
| Set 10 | 86(83) | **35** | 37 | 39 | 36 |
| **Sum** | 433 (403) | **120** | 125 | 133 | 129 |

**Table s3.** Distance error (deviation from additivity) for NSB-TK4 and Jellyfish-JC on bacterial dataset.

|  | # taxa (branches) | TotalFM (NSB-TK4) | totalFM (Jellyfish-JC) | TotalOLS (NSB-TK4) | TotalOLS (Jellyfish-JC) |
| --- | --- | --- | --- | --- | --- |
| Set 1 | 34(31) | **0.8209** | 1.29 | **0.032** | 0.0436 |
| Set 2 | 43(40) | **0.8032** | 0.8686 | **0.0114** | 0.0121 |
| Set 3 | 32(29) | **0.2758** | 0.3534 | **0.0132** | 0.0139 |
| Set 4 | 38(35) | **1.7227** | 1.926 | 0.1134 | **0.094** |
| Set 5 | 43(40) | **2.3761** | 2.5785 | 0.1688 | **0.1367** |
| Set 6 | 46(40) | **2.7291** | 2.9287 | 0.2003 | **0.1633** |
| Set 7 | 34(31) | **0.6379** | 0.8103 | **0.0309** | 0.0323 |
| Set 8 | 41(38) | **1.8973** | 2.0946 | 0.1186 | **0.0893** |
| Set 9 | 39(36) | **0.9773** | 1.2537 | **0.0478** | 0.0494 |
| Set 10 | 86(83) | **42.0776** | 55.9276 | **3.4154** | 4.0831 |
| **Sum** | 433 (403) | **54.3179** | 70.0314 | **4.1518** | 4.7177 |

**Table s4.** GenBank accession numbers of yeast species/strains.

| Species/Strains | GenBank accession |
| --- | --- |
| *Saccharomyces arboricola* | GCF_000292725.1 |
| *Saccharomyces cerevisiae* | GCF_000146045.2 |
| *Saccharomyces eubayanus* | GCF_001298625.1 |
| *Saccharomyces jurei* | GCA_900290405.1 |
| *Saccharomyces kudriavzevii* | GCA_900682665.1 |
| *Saccharomyces mikatae* | GCA_000167055.1 |
| *Saccharomyces paradoxus* | GCA_002079145.1 |
| *Saccharomyces uvarum* | GCA_002242645.1 |

## Footnotes

1 Note, however, that the original paper [28] has a mistake and has the term (*S*_1_ + *Q*_1_) in (2) instead of (*S*_1_ − *Q*_1_). Substituting the values of *X*_(*T*) and *Y*_(*T*), as defined in Eqn. 2 of the original paper, to Eqn. 18 in the original paper results in (*S*_1_ − *Q*_1_) instead of (*S*_1_ + *Q*_1_).

## References

[1] M. Höhl and M. A. Ragan, “Is multiple-sequence alignment required for accurate inference of phylogeny?” Systematic Biology, vol. 56, no. 2, pp. 206–221, 2007.

[2] G. A. Wu, S.-R. Jun, G. E. Sims, and S.-H. Kim, “Whole-proteome phylogeny of large dsDNA virus families by an alignment-free method,” Proceedings of the National Academy of Sciences, vol. 106, no. 31, pp. 12 826–12 831, 2009.

[3] S.-R. Jun, G. E. Sims, G. A. Wu, and S.-H. Kim, “Whole-proteome phylogeny of prokaryotes by feature frequency profiles: An alignment-free method with optimal feature resolution,” Proceedings of the National Academy of Sciences, vol. 107, no. 1, pp. 133–138, 2010.

[4] C. Daskalakis and S. Roch, “Alignment-free phylogenetic reconstruction: Sample complexity via a branching process analysis,” Annals of Applied Probability, vol. 23, no. 2, pp. 693–721, 2013.

[5] B. Haubold, “Alignment-free phylogenetics and population genetics,” Briefings in Bioinformatics, vol. 15, no. 3, pp. 407–418, 5 2014.

[6] M. Bogusz and S. Whelan, “Phylogenetic Tree Estimation With and Without Alignment: New Distance Methods and Benchmarking,” Systematic Biology, vol. 66, no. 2, pp. 218–231, 2016.

[7] C.-A. Leimeister, S. Sohrabi-Jahromi, and B. Morgenstern, “Fast and accurate phylogeny reconstruction using filtered spaced-word matches,” Bioinformatics, vol. 33, no. 7, p. btw776, 1 2017.

[8] H. Yi and L. Jin, “Co-phylog: an assembly-free phylogenomic approach for closely related organisms.” Nucleic acids research, vol. 41, no. 7, p. e75, 4 2013.

[9] H. Fan, A. R. Ives, Y. Surget-Groba, and C. H. Cannon, “An assembly and alignment-free method of phylogeny reconstruction from next-generation sequencing data,” BMC Genomics, vol. 16, no. 1, p. 522, 12 2015.

[10] E. S. Allman, J. A. Rhodes, and S. Sullivant, “Statistically Consistent k -mer Methods for Phylogenetic Tree Reconstruction,” Journal of Computational Biology, vol. 24, no. 2, pp. 153–171, 2 2017.

[11] S. Sarmashghi, K. Bohmann, M. T. P. Gilbert, V. Bafna, and S. Mirarab, “Skmer: assembly-free and alignment-free sample identification using genome skims,” Genome Biology, vol. 20, no. 1, p. 34, 12 2019.

[12] B. Linard, K. M. Swenson, and F. Pardi, “Rapid alignment-free phylogenetic identification of metagenomic sequences,” Bioinformatics, vol. 35, no. 18, pp. 3303–3312, 2019.

[13] A. Zielezinski, S. Vinga, J. Almeida, and W. M. Karlowski, “Alignment-free sequence comparison: Benefits, applications, and tools,” 2017.

[14] T. H. Ogdenw and M. S. Rosenberg, “Multiple sequence alignment accuracy and phylogenetic inference.” Systematic biology, vol. 55, no. 2, pp. 314–328, 2006.

[15] G. Lunter, A. Rocco, N. Mimouni, A. Heger, A. Caldeira, and J. Hein, “Uncertainty in homology inferences: assessing and improving genomic sequence alignment.” Genome research, vol. 18, no. 2, pp. 298–309, 2 2008.

[16] Li-San Wang, J. Leebens-Mack, P. K. Wall, K. Beckmann, C. W. de Pamphilis, and T. Warnow, “The Impact of Multiple Protein Sequence Alignment on Phylogenetic Estimation,” IEEE/ACM Transactions on Computational Biology and Bioinformatics, vol. 8, no. 4, pp. 1108–1119, 7 2011.

[17] K. Bohmann, S. Mirarab, V. Bafna, and M. T. P. Gilbert, “Beyond DNA barcoding: The unrealized potential of genome skim data in sample identification,” Molecular Ecology, vol. 29, no. 14, pp. 2521–2534, 7 2020.

[18] M. V. Westbury, K. F. Thompson, M. Louis, A. A. Cabrera, M. Skovrind, J. A. S. Castruita, R. Constantine, J. R. Stevens, and E. D. Lorenzen, “Ocean-wide genomic variation in Gray’s beaked whales, Mesoplodon grayi,” Royal Society Open Science, vol. 8, no. 3, p. rsos.201788, 3 2021.

[19] D. Lang, M. Tang, J. Hu, and X. Zhou, “Genome-skimming provides accurate quantification for pollen mixtures,” Molecular Ecology Resources, vol. 19, no. 6, pp. 1433–1446, 11 2019.

[20] M. Balaban, S. Sarmashghi, and S. Mirarab, “APPLES: Scalable Distance-Based Phylogenetic Placement with or without Alignments,” Systematic Biology, vol. 69, no. 3, pp. 566–578, 5 2020.

[21] M. Balaban and S. Mirarab, “Phylogenetic double placement of mixed samples,” Bioinformatics, vol. 36, no. Supplement_1, pp. i335–i343, 7 2020.

[22] K. Tang, J. Ren, and F. Sun, “Afann: bias adjustment for alignment-free sequence comparison based on sequencing data using neural network regression,” Genome Biology, vol. 20, no. 1, p. 266, 12 2019.

[23] A.-K. Lau, S. Dörrer, C.-A. Leimeister, C. Bleidorn, and B. Morgenstern, “Read-SpaM: assembly-free and alignment-free comparison of bacterial genomes with low sequencing coverage,” BMC Bioinformatics, vol. 20, no. S20, p. 638, 12 2019.

[24] T. H. Jukes and C. R. Cantor, “Evolution of protein molecules,” in Mammalian protein metabolism, Vol. III (1969), pp. 21–132, 1969, vol. III, pp. 21-132.

[25] S. Tavaré, “Line-of-descent and genealogical processes, and their applications in population genetics models,” Theoretical Population Biology, vol. 26, no. 2, pp. 119–164, 10 1984.

[26] O. Zagordi and J. R. Lobry, “Forcing reversibility in the no-strand-bias substitution model allows for the theoretical and practical identifiability of its 5 parameters from pairwise DNA sequence comparisons,” Gene, vol. 347, no. 2 SPEC. ISS., pp. 175–182, 2005.

[27] M. Steel, “Recovering a tree from the leaf colourations it generates under a Markov model,” Applied Mathematics Letters, 1994.

[28] N. Takahata and M. Kimura, “A model of evolutionary base substitutions and its application with special reference to rapid change of pseudogenes,” Genetics, 1981.

[29] S. Röhling, T. Dencker, and B. Morgenstern, “The number of k-mer matches between two DNA sequences as a function of k,” bioRxiv, p. 527515, 2019.

[30] K. Tamura and M. Nei, “Estimation of the Number of Nucleotide Substitutions in the Control Region of Mitochondrial-DNA in Humans and Chimpanzees,” Molecular biology and evolution, vol. 10, no. 3, pp. 512–526, 1993.

[31] M. Hasegawa, H. Kishino, and T. a. Yano, “Dating of the human-ape splitting by a molecular clock of mitochondrial DNA,” Journal of Molecular Evolution, 1985.

[32] N. Sueoka, “Intrastrand parity rules of DNA base composition and usage biases of synonymous codons,” Journal of Molecular Evolution, vol. 40, no. 3, pp. 318–325, 1995.

[33] C. I. Wu and N. Maeda, “Inequality in mutation rates of the two strands of DNA,” Nature, 1987.

[34] J. R. Lobry, “Properties of a general model of DNA evolution under no-strand-bias conditions,” Journal of Molecular Evolution, vol. 40, no. 3, pp. 326–330, 1995.

[35] B. D. Ondov, T. J. Treangen, P. Melsted, A. B. Mallonee, N. H. Bergman, S. Koren, and A. M. Phillippy, “Mash: fast genome and metagenome distance estimation using MinHash,” Genome Biology, vol. 17, no. 1, p. 132, 12 2016.

[36] C. Jain, L. M. Rodriguez-R, A. M. Phillippy, K. T. Konstantinidis, and S. Aluru, “High throughput ANI analysis of 90K prokaryotic genomes reveals clear species boundaries,” Nature Communications, vol. 9, no. 1, p. 5114, 12 2018.

[37] G. Marçais and C. Kingsford, “A fast, lock-free approach for efficient parallel counting of occurrences of k-mers,” Bioinformatics, 2011.

[38] Q. Zhu, U. Mai, W. Pfeiffer, S. Janssen, F. Asnicar, J. G. Sanders, P. Belda-Ferre, G. A. Al-Ghalith, E. Kopylova, D. McDonald, T. Kosciolek, J. B. Yin, S. Huang, N. Salam, J.-Y. Jiao, Z. Wu, Z. Z. Xu, K. Cantrell, Y. Yang, E. Sayyari, M. Rabiee, J. T. Morton, S. Podell, D. Knights, W.-J. Li, C. Huttenhower, N. Segata, L. Smarr, S. Mirarab, and R. Knight, “Phylogenomics of 10,575 genomes reveals evolutionary proximity between domains Bacteria and Archaea,” Nature Communications, vol. 10, no. 1, p. 5477, 12 2019.

[39] D. Robinson and L. Foulds, “Comparison of phylogenetic trees,” Mathematical Biosciences, vol. 53, no. 1-2, pp. 131–147, 1981.

[40] A. Bhattacharjee and M. S. Bayzid, “Machine learning based imputation techniques for estimating phylogenetic trees from incomplete distance matrices,” BMC Genomics, vol. 21, no. 1, p. 497, 12 2020.

[41] W. M. Fitch and E. Margoliash, “Construction of Phylogenetic Trees,” Science, vol. 155, no. 3760, pp. 279–284, 1 1967.

[42] Z. Yang and A. D. Yoder, “Estimation of the Transition/Transversion Rate Bias and Species Sampling,” Journal of Molecular Evolution, vol. 48, no. 3, pp. 274–283, 3 1999.

[43] W. Shen, S. Le, Y. Li, and F. Hu, “SeqKit: A cross-platform and ultrafast toolkit for FASTA/Q file manipulation,” PLoS ONE, vol. 11, no. 10, pp. 1–10, 2016.

[44] V. Lefort, R. Desper, and O. Gascuel, “FastME 2.0: A comprehensive, accurate, and fast distance-based phylogeny inference program,” Molecular Biology and Evolution, vol. 32, no. 10, pp. 2798–2800, 2015.

[45] M. Nei and T. Gojobori, “Simple methods for estimating the numbers of synonymous and nonsynonymous nucleotide substitutions.” Molecular biology and evolution, vol. 3, no. 5, pp. 418–26, 9 1986.

[46] L. Pritchard, R. H. Glover, S. Humphris, J. G. Elphinstone, and I. K. Toth, “Genomics and taxonomy in diagnostics for food security: soft-rotting enterobacterial plant pathogens,” Analytical Methods, vol. 8, no. 1, pp. 12–24, 2016.

[47] R. Chikhi and P. Medvedev, “Informed and automated k-mer size selection for genome assembly,” Bioinformatics, vol. 30, no. 1, pp. 31–37, 1 2014.

[48] D. N. Baker and B. Langmead, “Dashing: Fast and accurate genomic distances with HyperLogLog,” Genome Biology, 2019.

[49] A. Rahman, R. Chikhi, and P. Medvedev, “Disk compression of k-mer sets,” Algorithms for Molecular Biology, vol. 16, no. 1, p. 10, 12 2021.

